# Indigenous Reduction of Multi-drug Resistant Bacterial Pathogens in a Natural Uncontrolled Fermented Milk Ecosystem

**DOI:** 10.1101/2023.02.08.527581

**Authors:** Almira Akram, Muhammad Nadeem Khan, Waqas Ahmad, Shabir Ahmad Khan, Muhammad Imran

**Author notes:** **Corresponding author:** Dr. Muhammad Imran, Professor, Department of Microbiology, Faculty of Biological Sciences, Quaid-I-Azam University Islamabad, Pakistan.; **Email:**.

## Abstract

Raw milk and its products have been questioned for microbiological safety. However, Dahi: an artisanally fermented milk product that is yet microbiologically unexplored is popularly known for therapeutic uses in public. Aiming safety and therapeutic assessment of dahi, samples of raw milk and dahi were analysed for microbiology and bacterial pathogens. The pathogens were also evaluated for antibiotic resistance. Further dahi samples were assessed for antipathogen effect. The total bacterial count of raw milk was between 3.80 × 10^02^ to 2.97 × 10^07^ and total fungal count was 2.50 × 10^01^ to 4.90 × 10^02^ whereas total bacterial and fungal count of Dahi was 3.00 × 10^01^ to 4.05 × 10^09^ and 2.00 × 10^01^ to 5.30 × 10^07^. Bacterial population of the Dahi was found dominated by lactic acid bacteria. In milk samples, *E. coli* was detected as 38%, *S. aureus* 28%, *L. monocytogenes* 3%, *Salmonella* 42% and *Pseudomonas spp* 31%. In Dahi samples, their percentages were 7%, 10%, 7%, 8% and 2% respectively. The detected pathogens were found resistant against different antibiotics especially to third and fourth generations of cephalosporin and other beta-lactam drugs. When assessed invitro, the low incidence of pathogens in the Dahi samples were associated to its inhibitory effect against pathogens. Dahi samples also inhibited the growth of antibiotic resistance ATCC strains. The inhibitory activity of the Dahi is due to the changes occurs during fermentation but not corelated to the pH of the Dahi.

## Introduction

Raw milk is still commonly used in most parts of the world to traditionally ferment milk into different products. The evidence for health benefits and consumer preference due to diversity in the taste of these products are the established facts. Due to minimal or no pre-processing of milk, pathogen could be transferred to these products (Gonzales-Barron *et al*., 2017). Traditionally fermented raw milk products have been associated with complex microbial communities contributing to sensorial attributes and safety of the products (Bengoa *et al*., 2019). Therefore, the benefits of the microbes present in dairy and their role in the development of organoleptic properties because of fermentation are undeniable. Whereas at one side, dairy products contain ample amount of probiotic strains that have a lot of health benefits while on the other side, they may also harbor pathogenic microbes which lead to the spoilage of products ultimately causing diseases (Gonzales-Barron *et al*., 2017). The increased consciousness of consumers about safety of dairy products demands for safe products that are even free from traces of pathogens (Castellano *et al*., 2017; Newell *et al*., 2010; Schlundt, 2002). Moreover, the influence of microbes in food matrices directly impact the composition of the gut microbiota, clearly signifies the concerns about food-safety (González *et al*., 2019). Milk provides the best medium for the growth of several spoilage and pathogenic microbes that result in quality deterioration and ultimately cause infections in the consumer (Frank, 2009). Naturally, milk is believed to be sterile inside the udder of healthy farm animal, but it gets contaminated by handling procedures throughout the supply chain (Derakhshani *et al*., 2020). Milk quality varies rapidly due to direct or indirect contact with several microbial sources at the dairy farm and during transportation chain. This may include air, soil, animal dung, water, feeding material, udder surface, milking utensils and hygienic condition of workers at farm and storage conditions, cleanliness of storage units, duration and temperature of processing units, packaging and post-processing packaging of the product during supply chain (Angulo *et al*., 2009; Mallet *et al*., 2012; Michel *et al*., 2001; Oikonomou *et al*., 2020; Verdier-Metz *et al*., 2009). Numerous outbreaks of *Salmonella* spp, *L. monocytogenes* and O157, Shiga toxin producing *E. coli* have been reported due to consumption of unprocessed milk, signifying a great threat to the community health all over the world (Cdc, 2001, 2003; Mazurek *et al*., 2004; Msalya, 2017; Proctor & Davis, 2000; Reed & Grivetti, 2000).

Among fermented milk products, Dahi is the oldest and a highly consumed dairy product in Pakistan, produced by using thermally processed milk that has been given a sub-pasteurization temperature followed by addition of remaining junk of mature Dahi (Masud, 1991). Dahi on one side, is an important part of daily diet, and on the other hand it is popular in general population due to its therapeutic properties (Ahmad *et al*., 2012). Dahi is a nutritious food containing almost all the nutrients that are present in milk, but in a more digestive form. As there are no well described criteria for fermenting milk into Dahi, thus its quality is highly variable and can’t be considered analog to yogurt regarding its microbiology and sensory attributes. It has been reported that Dahi mainly carry bacteria species of *Lactobacillus, Lactococcus* and *Enterococcus* (Nawaz *et al*., 2019). Dahi differs from yogurt in respect of undefined starter culture and conditions of manufacturing (milk quality, culture concentration, viability, incubation time and temperature etc.). while is equivalent to yoghurt in terms of color, taste and texture (Robinson & Tamime, 2006). Studies have been elaborated the antimicrobial potential of different lactic acid bacteria isolated from the Dahi (Javed *et al*., 2011; Maqsood *et al*., 2013; Maqsood *et al*., 2008). It has been proposed that antipathogenic activity of Dahi is quite high because of which less pathogenic microorganisms survive in it. Its antagonistic potential has been reported against *Escherichia coli, Bacillus subtilis, Staphylococcus aureus* (Gandhi & Nambudripad, 1901) *Bacillus cereus* and *Salmonella typhosa* (KUMAR, Solanky, & Chauhan, 2003). Low pH, presence of bacteriophages, and certain compounds like lactic acid, acetic acid, ethanol, aromatic compounds, bacteriocins, exopolysaccharides and enzymes produced by LAB can act as antipathogen agents (Vuyst, 2004). Due to the broad range of antimicrobial agents, it is quite possible that multi-drug resistant pathogens could not survive in the Dahi ecosystem.

In this context, the present study is designed to evaluate microbiological safety status of the raw milk used for Dahi production. And antagonistic effect of the Dahi; a natural and uncontrolled fermentation ecosystem against antibiotic resistant pathogens.

## 2. Material and Methods

### 2.1 Sample collection

Samples were collected at two different stages from cottage scale dairies in twin cities of Rawalpindi and Islamabad, Pakistan. Initially 43 samples of Raw milk and Dahi samples were collected and at second stage 100 Dahi samples were collected. Samples were collected aseptically in sterile containers and transferred to the laboratory within one hour of sampling and processed immediately. All the samples were subjected to pH measurement and lactic acid content determination.

### 2.2. Microbiological analysis of Raw Milk and Dahi

The samples were analyzed for Total Aerobic Plate Counts (TAPC), Total yeast/mold count, *Lactobacilli, Lactococcal, Streptococci, Enterococci* and Gram Negatives count through spread plate techniques (Quinn *et al*., 2002). After homogenization, 1 ml of the raw milk/1 g of Dahi was homogenized with 20 ml of plate count agar and poured into sterile petri plate (PCA; Oxide CM0463 UK) and Oxytetracycline glucose agar (OGA) for enumeration of bacteria and fungi respectively. For enumeration of bacteria, plates were incubated for 24 h at 37°C and for 48 h at 30°C while for 5 days at same temperature for enumeration of yeasts/molds. Assumed *Lactobacilli* were enumerated on de Man Rogosa Sharpe (MRSA, CM0359, Oxoid) adjusted to pH 5.2 after incubating at 42°C for 72 h. The M17 (Oxoid) media was used for *Lactococci* after incubating plates at 37°C for 48 h. MacConkey agar (CM0005, Oxoid) was used for Enterobacteria and coliforms after incubation of plates at 37°C for 72 h.

### 2.3. Detection of pathogens in Raw Milk and Dahi

Selected pathogens *L. monocytogenes, E. coli, S. aureus, Salmonella* spp. and *Pseudomonas* spp. were detected using respective media. The detection of pathogens was done by direct plating the pathogens on respective media and then incubating at 37°C for 48 h. The *L. monocytogenes* was plated on Oxford agar (Liofilchem, Italy), *E. coli, S. aureus, Salmonella* spp and *Pseudomonas* spp on eosin methylene blue agar (EMB agar), mannitol salt agar (MSA), mannitol lysine crystal violet brilliant green (MLCB) agar and *Pseudomonas* Cetramide agar (PCA) respectively. Along with biochemical characterization, the identification of pathogens was confirmed through PCR by using specific primers.

Pathogenic bacteria in the samples were also detected by cultural independent method. For this purpose, specific PCR primers for each strain were either selected or developed. NCBI primer blast program http://www.ncbi.nlm.nih.gov/tools/primer-blast was used to generate the primers and to test *in silico* the specificity of the target for microorganisms. The specific PCR primers used in this study were manufactured by alpha DNA; Canada (**S. Table No. 1**).

DNA was extracted directly from milk and Dahi samples using phenol-chloroform method with few modifications for dairy products (Delbes *et al*., 2000). Subsequently, 2 ml of sample was taken in a micro-tube and centrifuged at 14000 rpm for 5 m. The supernatant was discarded, and the pellet was suspended in 400 µl of deionized water. 200 mg glass beads were added. Fifty microliter sodium dodecyl sulfate alkaline solution (Sigma-Aldrich) 10 mM EDTA (Sigma-Aldrich), 400 µL phenol–chloroform–isoamyl alcohol (Sigma-Aldrich) was vortexed for 2 minutes and cooled in ice bath for 3 minutes. The tubes were centrifuged at 14000 rpm for 30 minutes and upper aqueous phase was recovered after centrifugation and then two washing steps were performed first with phenol and chloroform iso-amyl alcohol (25:24:1) and second with chloroform isoamyl-alcohol (24:1). DNA was precipitated by addition of the double volume of absolute ethanol by incubating for 2 hours at −20ºC. Pellets were washed with 80% ethanol and dried. Final DNA was re-suspended in 100 µl of 1x Tris EDTA buffer and stored at 4°C until used. After extraction of DNA, polymerase chain reaction was performed for detection of the pathogens.

### 2.4. Antibiotic susceptibility assay of the detected pathogens

Antibiotic susceptibility of the isolated pathogens was determined against 23 antibiotics using Kirby Bauer disk diffusion method on Mueller-Hinton agar selected according to guidelines given by Clinical and Laboratory Standards Institute (CLSI-2021) recommended method. The following antibiotic susceptibility test disks: Gentamicin (CN) 10 µg, Ciprofloxacin (CIP) 5µg, Amoxicillin/clavulanic acid (AMC) 25 µg, Trimethoprim-sulfamethoxazole (SXT) 25 µg, Cefpirome (CPO) 30 µg, Meropenem (MEM) 10 µg, Ceftazidime (CAZ) 30 µg, Piperacillin-tazobactam (TZP) 110 µg, Amikacin (AK) 30 µg, Ceftriaxone (CRO) 30 µg, Imipenem (IPM) 10 µg, Ampicillin/Sulbactam (SAM) 20 µg, Amoxicillin/Clavulanate (AMC) 30 µg, Vancomycin (VA) 30 µg, Chloramphenicol (C) 30 µg, Teicoplanin (TEC) 30 µg, Ampicillin (AMP) 25 µg, Penicillin (P) 10 µg, Moxifloxacin (MXF) 5 µg, Fusidic Acid (FD) 10 µg, Cefazolin (KZ) 30 µg, Oxacilin (OX) 1 µg (Oxide-UK) were used.

### 2.5. Antipathogenic activity of Dahi

The indigenous antipathogenic effect of Dahi evaluated against the detected pathogens along with antibiotic resistance ATCC reference strains. Hundred Dahi samples that were collected in the second phase of Sample collection were used to determine antipathogenic activity. The strains that were used in the current study are *Listeria monocytogenes* ATCC 19115, *Staphylococcus aureus* ATCC 33592, *Salmonella enterica* subsp enterica ATCC 11013, *Bacillus subtilis* subsp. spizizenii ATCC 6633, *Escherichia coli* ATCC 1101362 and *Pseudomonas aeruginosa* ATCC BAA2108). Antipathogenic activity of Dahi was analyzed against each pathogen by using a well diffusion assay. The agar overlay method was used for formation of lawn. Zone formation was checked after 24 h.

### 2.6. Physiochemical properties of Dahi

Fourier Transform Infrared Spectroscopy was performed for all the samples to have a guess of their physiochemical properties. FTIR machine by BRUKER Company, model Tensor 27 was used for this purpose. All the samples were centrifuged at 5000 rpm for 10 minutes to remove the water content and then ran through the software to get results of physiochemical analysis.

### Statistical analysis

Microsoft Excel 2016 and GraphPad Prism 9 are used for statistical analysis. The data is presented mean of random duplicate or triplicate values.

## 3. Results

### 3.1. Raw milk and Dahi sampling

A total of forty-three raw Milk and Dahi samples were collected from cottage scale dairies for this study at the initial stage. And 100 Dahi samples were collected on the second stage. Among these, 100. 43 were the dairies from where samples were selected at initial stage. Remaining Dahi producers also used similar production methods for fermenting raw milk into Dahi.

### 3.2. General microbiology of raw milk and Dahi samples

The APC of raw milk was between 3.80 × 10^02^ to 2.97 × 10^07^ cfu/ml with an average of 1.58 × 10^06^ cfu/ml. A very low number of raw milk samples were found positive for yeast/fungal with very low counts. MRS media assumed *Lactobacilli* ranged from 3.21 ×10^4^ cfu/ml to 1.0 ×10^8^ cfu/ml with average count of 4.46 × 10^6^ cfu/ml. Cocci bacteria on M17 ranged from 5.29 × 10^4^ cfu/ml to 8.00 × 10^8^ cfu/ml with average count of 9.13 × 10^7^ cfu/ml. Gram negative count was maximum of 1.12 ×10^4^ cfu/ml while minimum was 1.00 × 10^1^ cfu/ml with an average of 3.60 × 10^3^ cfu/ml. The results for average microbial plate count are shown in (**Figure 1.A**). There were 11.6 % samples with high initial microbial count having TPC > 1, 000, 000 CFU/ml, 9.3% had moderate microbial count, between 500,001 and 1,000,000 CFU/ml and 44.10 % had low bacterial count TPC < 100,000 CFU/ml (**Figure 1.B**). The variation in bacterial load was not correlated (r = −0.34) with milk pH **(Table 1)**.

**Table 1:** Co-relation between APC and pH of the raw Milk

| pH | APC | Correlation |
| --- | --- | --- |
| 6.7 ± 0.18 | $1.58 \times 10^6 \pm 5.36 \times 10^6$ | -0.34 |

**Figure 1:**
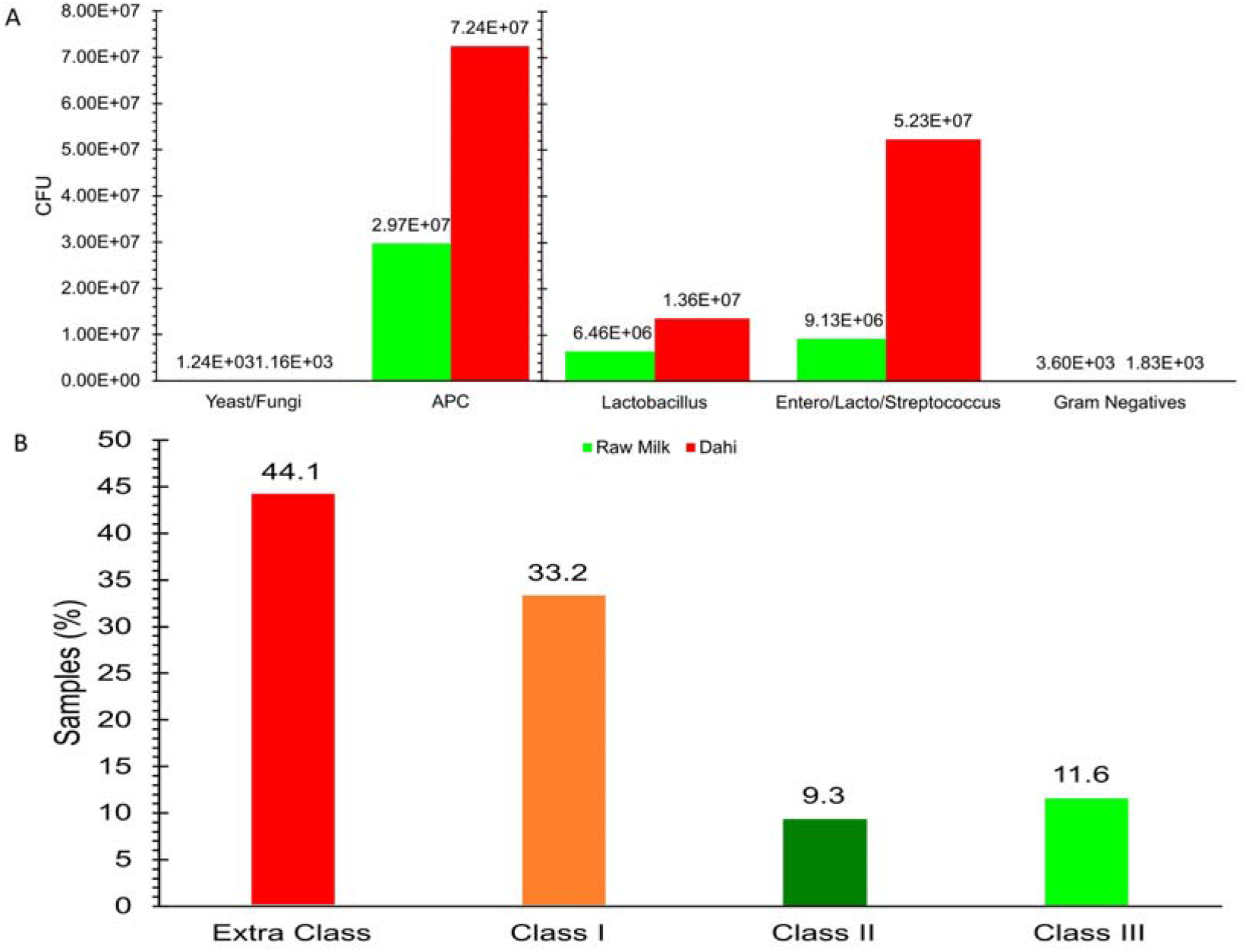
Microbiology of Raw Milk and Dahi samples. (A) Yeast, Aerobic plate, lactic acid bacteria, and Gram negatives count of Raw Milk and Dahi samples (B) Percentage of extra, I, II and III class of raw milk, Extra class raw milk: APC < 100 000 CFU/ml, I class raw milk: between 100, 001 and 500, 000 CFU/ml, II class raw milk: between 500, 001 and 1, 000, 000 CFU/ml and III class raw milk > 1, 000, 000 CFU/ml

For Dahi samples, the APC was between 3.00 × 10^01^ to 4.05 × 10^9^ CFU/gram of Dahi where assumed Lactic acid bacteria dominated the bacterial population. Yeas/fungi were also detected in Dahi samples. The average yeast/fungal count was above 1.16 × 10^03^ with minimum count of 1 × 10^02^ cfu/g and maximum 3.80 × 10^07^ cfu/g. On MRS media presumed Lactobacilli ranged between 2.15×10^4^ cfu/g to 1.21×10^8^ cfu/g with average count of 1.36× 10^7^ cfu/g. Cocci bacteria on M17 ranged from 6.75 × 10^4^ cfu/g to 5.51 × 10^8^ cfu/g with average count of 5.23 × 10^7^ cfu/g. Gram negative count was observed maximum as 2.62 × 10^4^ cfu/g and minimum as 1.50 × 10^1^ cfu/g with average count of 1.83 × 10^3^ cfu/g. Results for average microbial plate count are shown in (**Figure 1.A**).

### 3.3. Incidence of pathogens, and pattern of antibiotic resistance

When the samples of Milk and Dahi were analyzed for the detection of pathogens through culture dependent and culture independent methods. In Milk samples, the pathogens *E. coli, S. aureus, L. monocytogenes, Salmonella spp*. and *Pseudomonas spp*. were detected to be 38%, 28%, 4*%*, 46 % and 36 % respectively through culturing method. While by culture independent method, *E. coli, S. aureus, L. monocytogenes, Salmonella Spp*. and *Pseudomonas spp*. were detected as 30%, 18%, 12%, 20% and 16 % respectively. Whereas low incidence of pathogens was observed in Dahi samples. Ten samples (7%) were positive for both *Salmonella* spp and *E. coli*. Thirteen (9%) samples were positive for *S. aureus* and *Pseudomonas* spp while *L. monocytogenes* was detected only in one sample (0.69%) through culturing method. Ten samples (7%) were found positive for *E. coli* and three (7%) for *Salmonella* spp while no other pathogens were detected through culture independent method **(Figure 2A)**.

**Figure 2:**
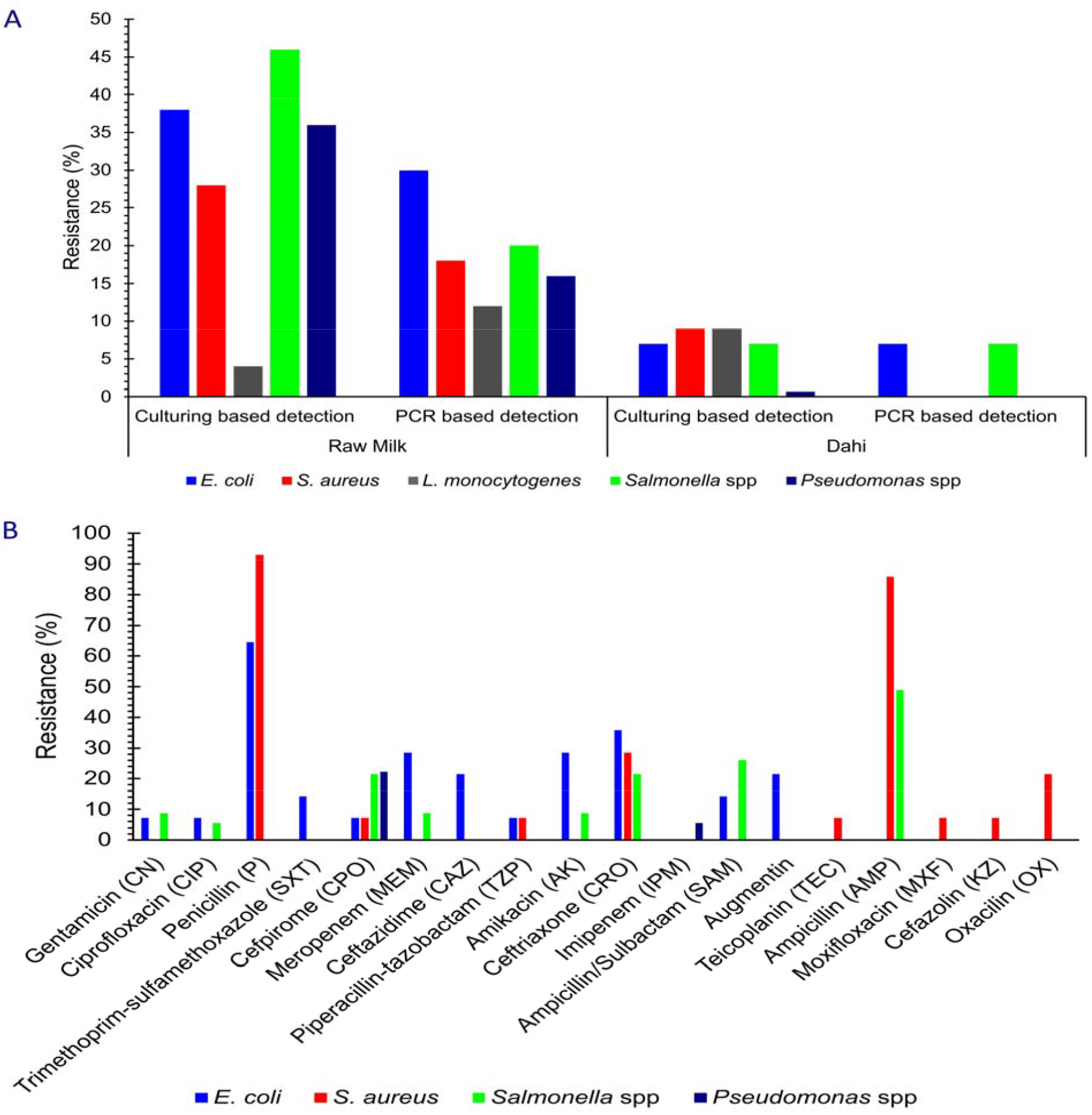
Pathogen incidence and antibiotic resistance of the detected pathogens (A) Comparative pathogen incidences in raw milk and Dahi samples (B) Antibiotic resistance pattern of the pathogen’s isolates including: *Salmonella* spp (n = 23), *Pseudomonas* spp (n = 18), *E. coli* (n = 19), *S. auras* (n = 14) and *L. monocytogenes* (n = 2)

When the pathogens isolated from raw milk were evaluated for their antibiotic resistance pattern. It was observed that that 69.6% of *Salmonella spp*. were resistant to Augmentin, 48.9 % to ampicillin, 21.8 % to ceftriaxone and cefpirome, 26.1 % to ampicillin with sulbactam, 8.7 % to gentamicin, amikacin and meropenem but showed no resistance to piperacillin/tazobactam. There were 92.86 % of *S. aureus* resistant to penicillin, 85.8 % to ampicillin, 50 % to fusidic acid, 28.6 % to ceftriaxone, 21.5% to oxacillin, 7.2 % to piperacillin/tazobactam, teicoplanin, moxifloxacin, cefpirome and cephazolin and showed no resistance to vancomycin, chloramphenicol, and trimethoprim sulfamethoxazole respectively. Among the isolated *E. coli* 64.5 % showed resistance to penicillin, 35.8% to ceftriaxone, 28.6 % to amikacin and meropenem, 21.5% to ceftazidime and augmentin, 14.3 % to ampicillin with sulbactam and trimethoprim sulfamethoxazole, 7.2 % to gentamicin, ciprofloxacin, and cefpirome and piperacillin/tazobactam and showed no resistance to imipenem. It was observed that 33.4 % of *Pseudomonas spp* were resistant to trimethoprim/sulfamethoxazole, 22.3 % to cefpirome, 5.6 % to imipenem and ciprofloxacin and showed no resistance to gentamicin, piperacillin/tazobactam, amikacin and ceftazidime respectively **(Figure 2B)**.

### 3.4. Antipathogenic activity of Dahi

Antipathogenic activity of the Dahi was checked through zone measurement. Larger zones showed strong antimicrobial activity and smaller zones showed weak antimicrobial activity. Antimicrobial activity of each sample was found to be different against each pathogen. Highest activity was shown against *Listeria monocytogenes* whereas lowest activity was observed against *Pseudomonas aeruginosa*. Among the total samples tested, 87% of the samples showed activity against *Listeria monocytogenes* (**Figure 3.A**). The largest zone of inhibition against *Listeria* was measured to be 31 mm whereas smallest was 1 mm with mean zone of 4.9 mm (**Figure 3.B)**. About 63% of Dahi samples showed positive results against *E. coli* (**Figure 3.A**). Largest zone of inhibition against *E. coli* was observed as 12 (**Figure 3.B**). Fifty-two percent of samples showed positive results against *S. enterica* and *B. subtilis* (**Figure 3.A**). Maximum zone of inhibition against *E. enterica* and *B. subtilis* were observed as 21 mm and 25 mm respectively (**Figure 3.B**). Only 20% of Dahi samples showed antimicrobial activity against *P. aeruginosa* (**Figure 3.A**) with maximum zone of inhibition 15 mm (**Figure 3.B**). Out of all the collected samples 23% of samples showed positive results against *S. aureus* (**Figure 3.A**) where maximum zone of inhibition is recorded as 21 mm (**Figure 3.B**). Sample and pathogen-wise results are provided in (**S. Data 1**).

**Figure 3:**
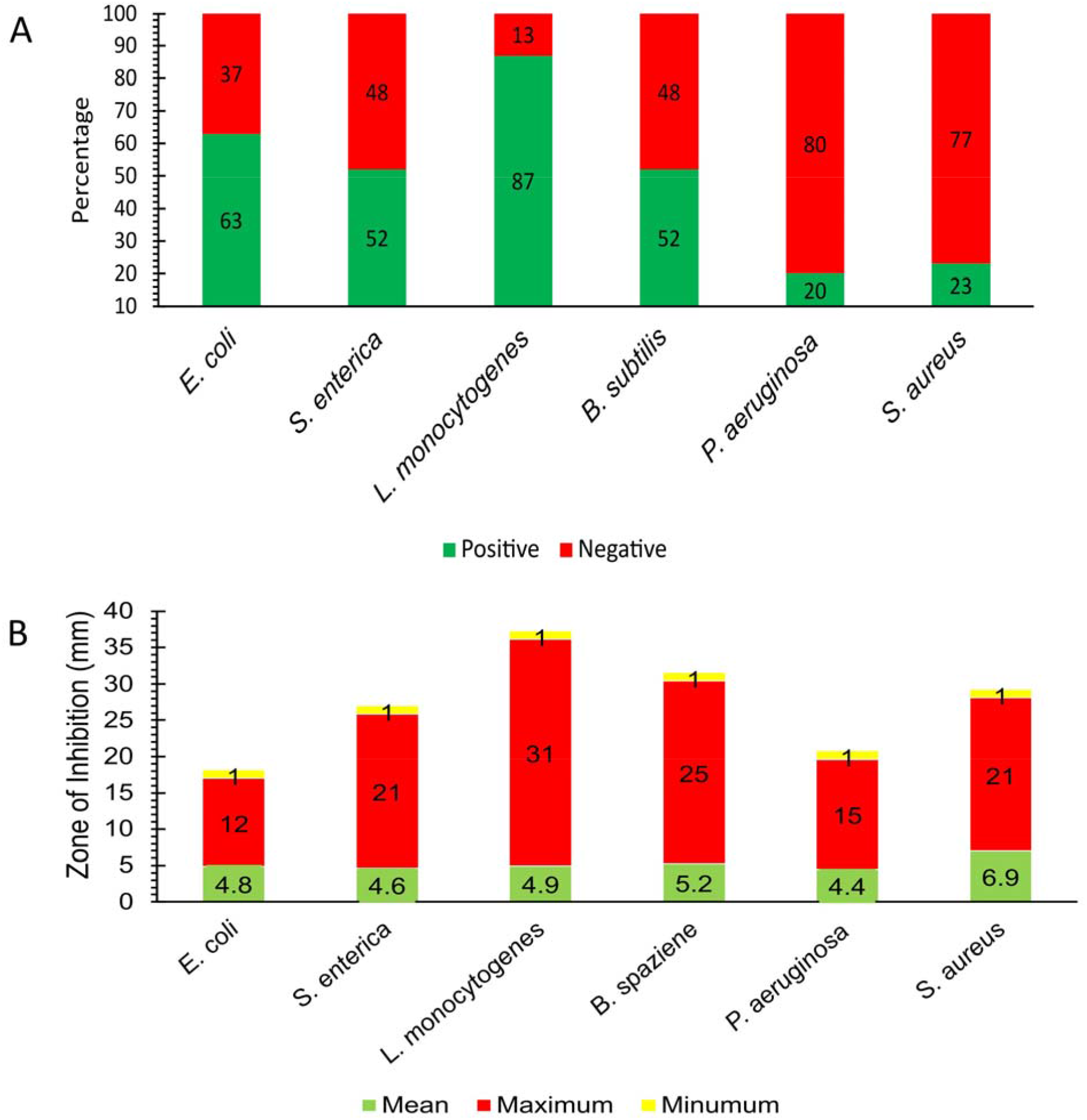
Antibacterial activity of Dahi samples (A) Number and percentage of Dahi samples positive and negative against selected pathogens (B) Zone of inhibition against selected pathogens.

### 3.5. Co-relation between pH and antimicrobial activity of Dahi

A correlation between pH and antimicrobial activity of Dahi samples was analyzed and it was observed that there is no co-relation between pH and antimicrobial activity of the Dahi samples (**Figure 4**).

**Figure 4:**
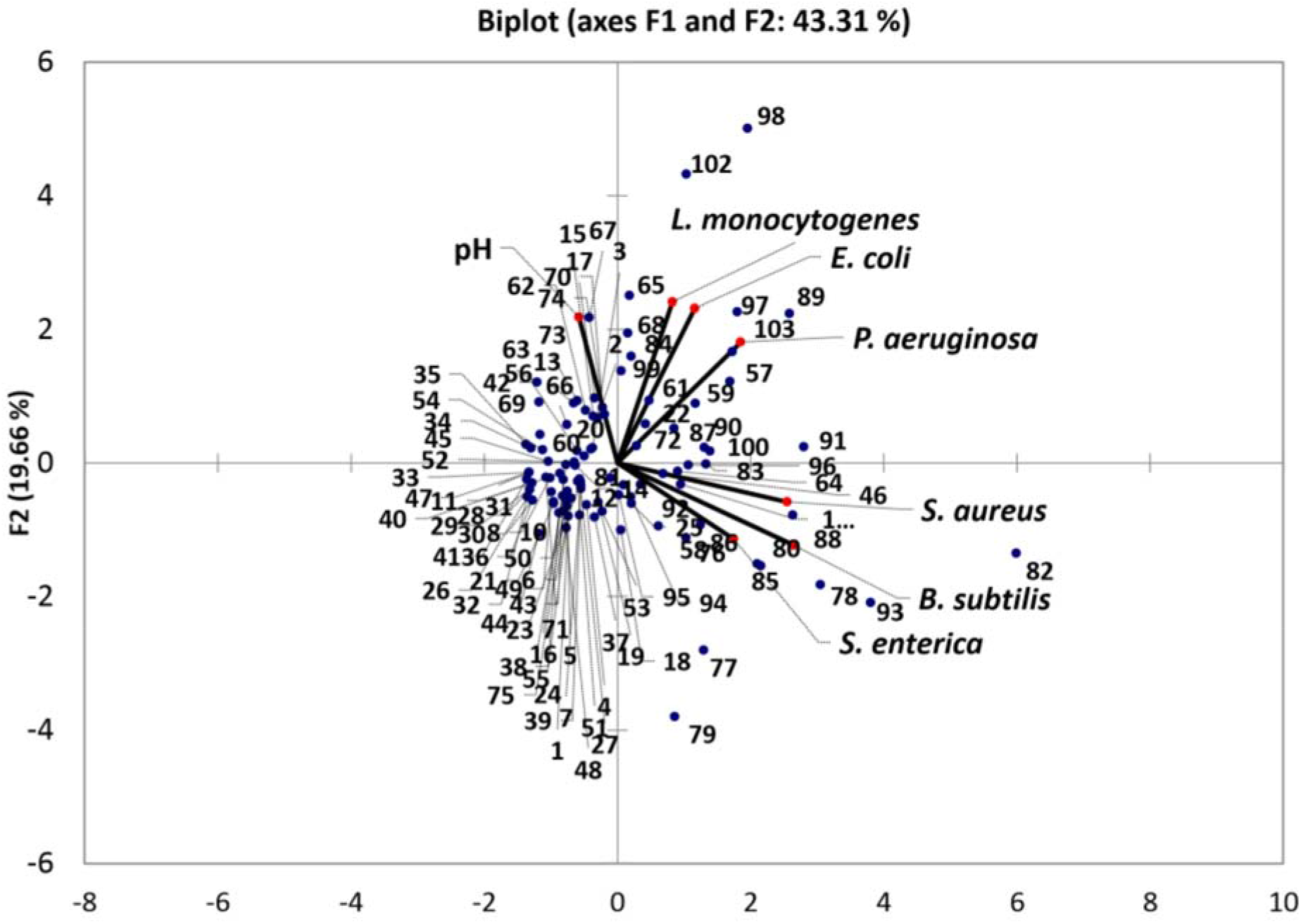
PCA Chart showing co-relation between pH and antimicrobial activity of the dahi samples.

### 3.6. Physiochemistry of the Dahi

The quality attributes and biochemical properties of yogurt are mainly due to the process of fermentation. These biochemical properties were observed by using the FTIR technique. In FTIR the position of functional group indicates the presence of a certain compounds (Jawhari *et al*., 1992). Major difference was observed in the physiochemistry of samples having good antimicrobial activity against pathogens and those lacking it. Samples showing high positive value were observed to have peaks at wave numbers 3593 cm^-1^ and 3543 cm^-1^ which indicate the presence of alcohols and phenols (**Figure 5**).

**Figure 5:**
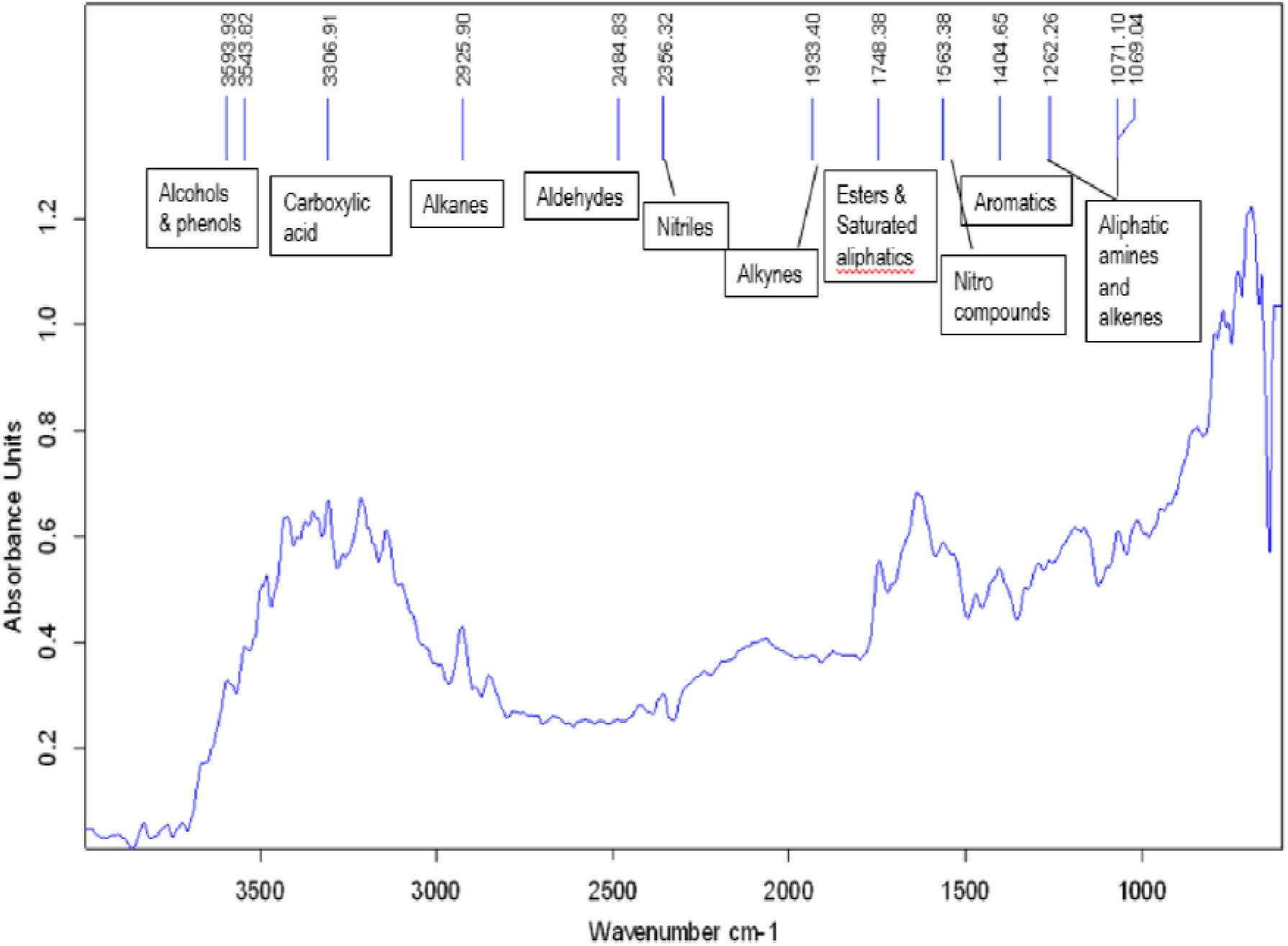
FTIR based physiochemistry of sample a representative Dahi sample having antimicrobial activity.

## Discussion

Traditionally fermented milk products have been consumed worldwide due to their unique organoleptic and therapeutic properties. The use of raw milk to produce these dairy products ha been associated with safety concerns of the products. The raw milk dairy products have been a focus of study due to safety challenges; however, they have higher consumption worldwide due to high nutritional values. However, in the countries where the dairy sector is not fully industrialized, production of these products is still challenging in terms of safety. Different outbreaks have been associated with consumption of raw milk (Gruber *et al*., 2020). Current study aimed to assess the microbiological profile of Raw Milk and the most consumed dairy fermented product Dahi. In the current study, raw milk and Dahi samples collected from cottage scale shops with different hygienic conditions, geographic locations, and preservation methods showed different level of microbial loads. In spite of poor hygienic conditions more than forty percent of milk samples had low bacterial count (<100000 cfu/ml) in comparison to many countries where farming systems are still conventional (Nádia *et al*., 2012). Lactic acid bacteria are present as dominant microflora in most of the Dahi samples which convert lactose to lactic acid thereby reducing survival of pathogenic bacteria (Garrote *et al*., 2000; Kingamkono *et al*., 1998). However, a number of pathogens can tolerate and survive unfavorable conditions created by lactic acid bacteria (Leyer *et al*., 1995). The presence of yeast and mold in milk can be related to the risk of production of toxins in Dahi. (Alborzi *et al*., 2006; Hussain & Anwar, 2008). While in Dahi Yeast play a synergistic role in milk fermentation that’s results in the thickening and souring of milk (Adesulu-Dahunsi *et al*., 2020).

The microbial load is an indicator of safety, in fermented products where it dependents on starter culture used, fermentation time followed and storage of the product (Tamine *et al*., 2007). In the present study, when pathogens were detected through culturing and PCR, the pathogens detected were similar, but their number was quite different for both the methods used. Both the methods are justified for their use where culturing can detect only viable bacteria while PCR can detect even the dead ones, that’s why the number detected by both the methods can be different (Saravanan *et al*., 2021). Milk samples of the current study were observed to have more *E. coli*. The contamination of *E. coli* O157 in raw milk occurs mainly through fecal contamination during milking but contamination of *E. coli* from infected udder during milking has also been reported (Hussein & Sakuma, 2005a; Lira *et al*., 2004). Thirty-eight percent of milk samples were positive for *E. coli* by culturing techniques while 30 % were positive using PCR. A study reported 58 % *E. coli* in soft and hard cheeses made from raw milk (Ansay & Kaspar, 1997). The incidence of *E. coli* was also reported in 20.8 % of Kareish cheese and 32.8 % in Damieth cheeses samples made from raw milk (Ombarak *et al*., 2016). The culture independent PCR based detection confirms only the result for *E. coli* from one sample. Dahi samples collected during current study were also observed to carry *E. coli*. But their number was low as compared to the raw milk. Acidic pH of yogurt, high temperature during its production and presence of probiotic lactic acid bacteria are among the known factors that attribute low prevalence of *E*.*coli* (Dehkordi, Yazdani, Mozafari, & Valizadeh, 2014Bachrouri, Quinto, & Mora, 2006). However, the ability of *E. coli* to resist acidic pH and low temperature could be the reason of its survival in the Dahi (Hsin-Yi & Chou, 2001; Morgan et al., 1993).

Four samples of raw milk were found positive for both *S. aureus* which could be related to mastitis in the milking animals. In many other studies, the range of incidence has been reported between 13% to safety, it is to be noted that *S. aureus* is an enterotoxin producer but to produce the sufficient amount of toxin that could cause human illness, the pathogen concentration must exceed 5 log/ml (Jay *et al*., 2005). Studies has reported the prevalence of *S. aureus* in bovine was 100% (Adesiyun *et al*., 1998; Chye *et al*., 2004; Ortolani *et al*., 2010). Other studies have also reported the presence of *S. aureus* as 47.3% and 98.8% in raw milk samples which is significantly higher than findings of the current study (Jakobsen *et al*., 2011). The *S. aureus* was also present in the Dahi samples, but their number was quite low. The ability of *S. aureus* to adapt to acidic environment could be the reason its presence in Dahi samples (Pazakova, Turek, & Laciakova, 1997).

Low incidence of *L. monocytogenes* (12%) by PCR and (4%) by culturing techniques in raw milk could be related to the absence of indoor housing activity during the summer season, a similar fact has been indicated in a recent study (Hill *et al*., 2012). Contamination of milk by *L. monocytogenes* could be related with the indoor housing of cattle as well as poorly made silage and poor hygiene (Husu *et al*., 1990; Sanaa *et al*., 1993). Van *et al*., 2004 reported that 6.5% samples were positive for *L. monocytogenes* in bulk tank milk for US dairies and determined the prevalence of *L. monocytogenes* in bulk milk samples from 474 herds in Washington, Oregon, and Idaho. In another study, there was 8% of *Listeria spp*. in Turkish food (Yavuz *et al*., 2012). In current study no *L. monocytogenes* was detected in Dahi samples. It has been proved that conditions in yogurt are unfavorable for the growth of *Listeria monocytogenes*, however, changes in pH and storage time can support its growth in yogurt environment for some time (Cirone et al., 2013, Rubin & Vaughan, 1979). Dahi has a mixed culture of different Lactic Acid Bacteria and co-occurrence of this cocktail of bacteria and the bacteriocins produced by these bacteria, can control or inhibit the growth of *Listeria monocytogenes* (SCHAACK & MARTH, 1988). However, the presence of *Listeria* in Dahi can be due to use of unhygienic practices during yogurt production, use of contaminated milk and psychotropic nature of the *Listeria* (Borges, Silva, & Teixeira, 2011; Szczawiński, Szczawińska, & Łobacz, 2014).

*Salmonella* was relatively higher in milk samples of the current study. The range of *Salmonella* in different studies remained between “not detection” to 8.9% (D’amico *et al*., 2008; Hill *et al*., 2011, 2012; Jayarao *et al*., 2006; Murinda *et al*., 2002; Steele *et al*., 1997). The prevalence of *Salmonella spp*. was found to be 5% of the tested milk samples in a recent study (Fotou *et al*., 2011) and the prevalence of *Salmonella spp*.in food products in northern China was 20.9 % in 2010 (Yan *et al*., 2010). The prevalence of *Salmonella* in dairy farms was 47 % (Addis *et al*., 2011). *Salmonella* spp were also detected in Dahi samples but in very low number. Heavily contaminated milk with *Salmonella enterica* when used for the production of yogurt can be a source of *Salmonella* infection (Szczawiński et al., 2014).

In the present study, thirty percent of raw milk samples were contaminated with *Pseudomonas* spp. In spoiled UHT milk, it was found that among Gram-negative Psychotropic bacteria *Pseudomonas spp* were 67% which symbolizes the presence of contaminated water, soil or equipment thus decreasing the shelf life of milk (Chen *et al*., 2011). The high level of psychotropic bacteria can also produce heat stable enzymes which remained stable during pasteurization and could be responsible for spoilage of milk and milk products (Hayes & Boor, 2001). The lowest antimicrobial activity was observed against *Pseudomonas aeruginosa* i.e., 22%. this result is in contrast to a study conducted on mice where lung infection by Pseudomonas was cleared to a great extent by using yogurt and *Lactobacillus Casei* (Alvarez, Herrero, Bru, & Perdigon, 2001). On the other hand certain antibacterial resistant strains of *Pseudomonas aeruginosa* have been isolated from different food products (Driscoll, Brody, & Kollef, 2007). As the process of fermentation proceeds several low molecular weight compounds are produced by lactic acid bacteria. These compounds include the formation of hydrogen peroxide, carbon dioxide, diacetyl, and some other compounds. Lesser count of pathogens in yogurt can be attributed to these compounds (Omemu & Faniran, 2011).

When the selected pathogens were evaluated for antibiotic resistance, it was observed that 64.3% of *E. coli* were resistant to penicillin, 35.8% to ceftriaxone, 28.6% to amikacin, meropenem and 14.3% to ampicillin/sulbactam and trimethoprim/sulfamethoxazole. The *E. coli* isolates from a cow with mastitis has been reported to be 98.4% ampicillin resistant, less than 20% carbenicillin, gentamicin, cephalothin, trimethoprim, and amikacin, 3.1% ceftriaxone, 22.5% cefuroxime and 97.7 % aztreonam resistant (Srinivasan *et al*., 2007). Another similar study reported that *E. coli* O157 isolated from bovine, caprine, and ovine were resistant to ampicillin while tetracycline was found to be most effective against all isolates followed by gentamicin and cefuroxime (Solomakos *et al*., 2009). A study found that 27.8 % *E. coli* isolated from mastitis are resistant to at least one antimicrobial agent and 20.1% isolates are resistant to more than one antimicrobial agents. Where 18.6 % of the isolates were found ampicillin resistant, 16.4% streptomycin resistant, 15.7% tetracycline resistant and 13.6 % sulphamethoxazole resistant while gentamicin and florfenicol were found effective against all the isolates (Suojala *et al*., 2011).

Oxacillin resistance of *S. aureus* was (21.4%) out of 14 isolates determined by Kirby Bauer disk diffusion methods. All Oxacillin resistant *S. aureus* were sensitive to vancomycin and trimethoprim/Sulfamethoxazole. A similar study reported 4.8% MRSA isolated from caprine mastitis using Kirby Bauer method (Zeki *et al*., 2012). Similarly, among 421 isolates of *S. aureus* from major food animals 6.6 % were reported as oxacillin resistant (Lee, 2003).

It was found that 52% of the isolated *Salmonella* are resistant to ampicillin. It has been reported that 19.4% and 100% of the isolated *Salmonella* spp were found resistant to ampicillin (Addis *et al*., 2011; Endrias & Cornelius, 2009). In current study, Ciprofloxacin was found effective antimicrobial agent against all the isolated pathogens and this is comparable with results reported by (Akinyemia *et al*., 2005). Trimethoprim/sulfamethaxazole was very effective against all isolates of *Salmonella* species this is different from the finding as described by (Yan *et al*., 2010). *Salmonella* has shown 78.3% resistance to the third generation of cephalosporin such as ceftriaxone and Yan *et al* (2010) reported that *Salmonella* spp. isolated from retail food showed less than 5 % resistance to the third generation of cephalosporins.

In a current study it was found that 33.4% of the isolated *Pseudomonas spp*. are resistant to Trimethoprim/sulfamethaxazole, 100% sensitive to gentamicin, piperacillin/tazobactam, amikacin and ceftazidime and 94.4 % sensitive to imipenem. Similar results have been reported that *pseudomonas spp*. showed 100% resistance to penicillin G and 28.1% sulphamethoxazole/trimethoprim and all isolates were 100% sensitive to gentamicin, ciprofloxacin, ceftazidime, imipenem, and amikacin (Arslan *et al*., 2011).

When Dahi samples were evaluated for in-vitro antipathogenic effect. It was observed that Dahi was able to inhibit the growth of different pathogens at various ranges. The antibacterial properties of Dahi can be due to different factors such as low pH (lactic acid) (Gao *et al*., 2019), bacteriocins (Agriopoulou *et al*., 2020), and production of different antimicrobial compounds (Sakandar & Zhang, 2021). In Dahi production, the lactic acid bacteria work in synergy where different strains of lactic acid bacteria with different properties and abilities, are combined in a mixture, they can complement each other and make the environment unfavourable for undesired bacteria (Adesulu-Dahunsi *et al*., 2020). When the correlation between pH and antipathogenic activity of Dahi was drawn. It was found that the pH of the Dahi is not corelated with its antipathogenic effect. Upon FTIR spectroscopy, it might be suggested that alcohol and phenols play a vital role in antimicrobial activity of Dahi. It can be concluded that minor changes in physiochemical properties of Dahi, which may be due to the different qualities of milk used for Dahi formation or other reasons can greatly affect the antimicrobial activity and health attributes of Dahi.

## Conclusion

From current study, it was concluded that milk contains pathogenic microbes which may cause food poisoning and gastrointestinal infections. The pathogenic bacteria include *L. monocytogenes, S. aureus, Salmonella spp*. and *E. coli*. Proteolytic and psychotropic bacteria are present after delivery to dairy plants. These harmful bacteria in raw milk will greatly affect people with weak immunity. Despite pasteurization heat stable enzymes secreted by psychotropic organisms, may spoil milk and milk products. If the process of pasteurization is delayed, psychotropic bacteria such as *Pseudomonas spp*. will proliferate and contaminate milk. For microbiological assessment of milk rapid, reliable, and sensitive identification system is very important. During past decades use, overuse or misuse of antibiotics rose to the development of antibiotic resistant bacteria. Whereas on the other hand, Dahi a fermented milk product is comparatively safe. The incidence of pathogens in the Dahi is very low. The reason of low pathogen presence in the Dahi is due changes that occurs during fermentation.

## Notes

### Competing Interest Statement

The authors have declared no competing interest.

